# *De novo* assembly and annotation of the eastern fence lizard (*Sceloporus undulatus*) transcriptome

**DOI:** 10.1101/136069

**Authors:** Mariana B. Grizante, Marc Tollis, Juan J. Rodriguez, Ofir Levy, Michael J. Angilletta, Kenro Kusumi

## Abstract

**Background:** The eastern fence lizard (*Sceloporus undulatus*) has been a model species for ecological and evolutionary research. Genomic and transcriptomic resources for this species would promote investigation of genetic mechanisms that underpin plastic responses to environmental stress, such as climate warming. Moreover, such resources would aid comparative studies of complex traits at the molecular level, such as the transition from oviparous to viviparous reproduction, which happened at least four times within *Sceloporus*.

**Findings:** A *de novo* transcriptome assembly for *Sceloporus undulatus*, Sund_v1.0, was generated using over 179 million Illumina reads obtained from three tissues (whole brain, skeletal muscle, and embryo) as well as previously reported liver sequences. The Sund_v1.0 assembly had an average contig length of 782 nucleotides and an E90N50 statistic of 2,550 nucleotides. Comparing *S. undulatus* transcripts with the benchmarking universal single-copy orthologs (BUSCO) for tetrapod species yielded 97.2% gene representation. A total of 13,422 protein-coding orthologs were identified in comparison to the genome of the green anole lizard, *Anolis carolinensis*, which is the closest related species with genomic data available.

**Conclusions:** The multi-tissue transcriptome of *S. undulatus* is the first for a member of the family Phrynosomatidae, offering an important resource to advance studies of adaptation in this species and genomic research in reptiles.

## Data description

### Context

Eastern fence lizards belong to a clade, the *Sceloporus undulatus* complex, which spans much of the United States and northern Mexico [1]. Because these lizards occupy a wide range of habitats and environmental conditions, *S. undulatus* has been a good model for studies of organismal ecology [2–4], population dynamics [5,6], and local adaptation [7–9]. In particular, embryos of oviparous *S. undulatus* are subjected to oscillations in nest temperature that are known to affect development [10–13], which could potentially be compensated for by egg-laying behavior in adult females [14,15]. Embryos of this species have a threshold for thermal tolerance at high temperatures and are thus susceptible to potential warming due to climate change [11,16]. Other species in the genus *Sceloporus* evolved either prolonged or complete retention of eggs in response to cold environments. In fact, viviparity has evolved in association with cooler climates at least four times within *Sceloporus* and another two times in the Phrynosomatidae [17], along with numerous physiological and morphological adaptations expected to accompany this convergent trait. Specifically, a congeneric species *(S. jarrovi)* displays specialized features in the placenta, although relying mostly on yolk nutrients during development (lecithotrophy) [18,19]. A comparative study of gene expression among *Sceloporus* species that differ in parity mode (oviparous vs. viviparous) would allow testing for convergence with the pregnant-specific gene expression profile described for viviparous lizard species from family Scincidae, whose development depend mostly on nutrients from the mother (matrotrophy) [20]. To begin to identify the genes for molecular studies of these processes, we have sequenced and annotated a *de novo* multi-tissue transcriptome for *Sceloporus undulatus*.

## Methods

### a) Sampling

Gravid females of *Sceloporus undulatus* were collected in Edgefield County, South Carolina (33.7°N, 82.0°W) and transported to Arizona State University. These animals were maintained under conditions described in previous publications [21,22], which were approved by the Institutional Animal Care and Use Committee (Protocol #14-1338R). Approximately two days after laying eggs, each lizard was euthanized by injecting sodium pentobarbital into the coelomic cavity. The whole brain and skeletal muscle samples were removed and placed in RNA-lysis buffer (mirVana miRNA Isolation Kit, Ambion) and flash-frozen. Additionally, three early-stage embryos from each clutch were dissected, pooled together, and homogenized in RNA-lysis buffer using the same protocol.

### b) Sequencing

Total RNA was isolated from three tissue samples (whole brain, skeletal muscle and embryos) from each individual using the mirVana miRNA Isolation Kit (Ambion) protocol. Samples were checked for quality on a 2100 Bioanalyzer (Agilent). One sample from each tissue was selected for RNA-Seq based on the highest RIN, with a cutoff of 8.0. For each selected sample, 3 μg of total RNA was sent to the University of Arizona Genetics Core (Tucson, AZ) for library preparation and with TruSeq v3 chemistry for a standard insert size. RNA samples were multiplexed and sequenced using an Illumina HiSEq 2000 to generate 100-bp paired-end reads. Publicly available raw Illumina RNA-Seq reads from S. *undulatus* liver [23] were added to our dataset. After removing adaptors, raw reads from the four tissues were evaluated using FastQC (http://www.bioinformatics.babraham.ac.uk/projects/fastqc/, v-0.11.5) and trimmed using Trimmomatic (v-0.32, [24]), filtering for quality score (≥Q20) and using HEADCROP:9 to minimize nucleotide bias. This procedure yielded 179,374,469 quality-filtered reads. **Table 1** summarizes read-pair counts from whole brain, skeletal muscle, whole embryos, and liver.

**Table 1.** Number of pairs and accessions numbers for *Sceloporus undulatus* sequence reads.

| Tissue | Number of read pairs | Accession numbers |
| --- | --- | --- |
| *This study* |  |  |
| Whole Brain | 51,537,265 | SAMN06312741 |
| Embryo | 49,112,293 | SAMN06312742 |
| Skeletal muscle | 42,922,488 | SAMN06312743 |
| *McGaugh et al., 2015* |  |  |
| Liver | 35,802,423 | SRR629640 |
| Total | 179,374,469 | — |

### c) Assembly and annotation

All trimmed reads were pooled and assembled *de novo* using Trinity (v-2.2.0, default k-mer size of 25 [25]), which is an efficient transcriptome assembly method for non-model species without a reference genome available [26]. The most comprehensive transcriptome, obtained using reads from four tissues, consists of 547,370 contigs with an average length of 781.5 nucleotides (**Table 2**)—shorter than other assemblies because of the range of contig sizes that varied among datasets (1, 3 and 4 tissues; **Table S1, Fig. S1**). The N50 of the most highly expressed transcripts that represent 90% of the total normalized expression data (E90N50) was highest in the assembly based on four tissues, hereafter referred to as Sund_v1.0 (**Table 2**). A subset of contigs containing the longest open reading frames (ORFs), representing 123,323 transcripts, was extracted from the Sund_v1.0 assembly using TransDecoder (v-3.0.0, http://transdecoder.github.io) with homology searches against the databases UniProtKB/SwissProt [27] and PFAM [28]. The transcriptome obtained was annotated using Trinotate (v-3.0, http://trinotate.github.io), which involved searching against multiple databases (as UniProtKB/SwissProt, PFAM, signalP, GO) to identify sequence homology and protein domains, as well as to predict signaling peptides. **Table 3** summarizes the annotation results.

**Table 2.** Statistics for the *de novo* assembly of *Sceloporus undulatus* transcriptome (Sund_v1.0).

| **Assembly** | **1 tissue [23]** | **3 tissues** | **4 tissues**<br>**(Sund_v1.0)** |
| --- | --- | --- | --- |
| Total of Trinity transcript contigs | 158,323 | 492,249 | 547,370 |
| Total of Trinity 'genes' | 138,031 | 422,687 | 467,658 |
| GC% | 43.8 | 42.9 | 42.8 |
| Contig N50 (bp) | 1,720 | 1,648 | 1,438 |
| Contig E90N50 (bp) | 2,254 | 2,640 | 2,550 |
| Average contig length (bp) | 833.0 | 822.4 | 781.5 |
| Transcripts with the longest | 86,630 | 212,172 | 217,756 |
| ORFs | (54.7%) | (43.1%) | (39.8%) |

The different assemblies used data from a previous study (1 tissue, liver [23]), data from this study (3 tissues:: whole brain, skeletal muscle, embryos), and the two datasets combined (4 tissues, or Sund\_v1.0).

**Table 3.** Annotation summary of *Sceloporus undulatus de novo* transcriptome assembly (Sund_v1.0).

| Annotation of the Sund_v1.0 assembly |  |
| --- | --- |
| Annotated genes | 467,658 |
| Annotated transcript isoforms | 547,370 |
| Annotated isoforms/gene | 1.17 |
| Transcripts with Swiss-Prot annotation | (71,944) |
| Transcripts with PFAM annotation | 51,018 (46,432) |
| Transcripts with KEGG annotation | 65,694 (21,520) |
| Transcripts with GO annotation | 73,936 (66,554) |

Unique annotation numbers are indicated by parentheses.

## Data validation and quality control

Trimmed reads were aligned back to the assembled contigs using Bowtie2 (v-2.2.6 [29]). From the 176,086,787 reads that aligned, 97% represented proper pairs (**Table S2**), indicating good read representation in the Sund_v1.0 assembly. To assess quality and completeness of the assemblies, we first compared the Sund_v1.0 transcripts with the BUSCO profile for Tetrapoda (BUSCO v-2.0 [30]), which has BLAST+ (v-2.2.31 [31]) and HMMER (v-3.1b2 [32]) as dependencies. This procedure revealed that the Sund_v1.0 assembly captured 97.1% of the expected orthologs, a result comparable to the 97.8% obtained for *Anolis carolinensis* transcriptome using 14 tissues [33] (**Table 4**). Next, nucleotide sequences of Sund_v1.0 transcripts with the longest ORFs were compared to the protein set of *Anolis carolinensis* (AnoCar2.0, Ensembl) using BLASTX (evalue=1e-20, max_target_seqs=1). This comparison showed that 11,223 transcripts of S. *undulatus* have nearly full-length (>80%) alignment coverage with *A. carolinensis* proteins (**Table S3**). Predicted proteins of S. *undulatus* were also used to identify 13,422 one-to-one orthologs with proteins of *A. carolinensis* through reciprocal BLAST (evalue=1e-6, max_target_seqs=1).

**Table 4.** BUSCO results for the transcriptomes of *Sceloporus undulatus* and *Anolis carolinensis*.

|  | *Sceloporus undulatus* |  |  | *Anolis carolinensis* |
| --- | --- | --- | --- | --- |
|  | 1 tissue | 3 tissues | 4 tissues<br>(Sund_v1.0) | 14 tissues |
| Complete genes | 72.5% | 91.7% | 92.3% | 96.7% |
| Duplicated genes | 25% | 43.8% | 43.9% | 37.9% |
| Fragmented genes | 9.2% | 4.8% | 4.8% | 1.1% |
| Missing genes | 18.3% | 3.5% | 2.9% | 2.2% |
| Reference | McGaugh et al, 2015 [23] | This study | This study | Eckalbar et al, 2013 [33] |

For *S. undulatus*, the Sund\_v1.0 assembly includes 4 tissues, specifically 3 tissues from this study (whole brain, skeletal muscle and embryos) and 1 previously reported tissue (liver [23]). For *A. carolinensis*, transcriptomes included adrenal gland, brain, dewlap skin, embryos, and pooled samples, heart, liver, lung, original tail, ovary, regenerating tail tip, regenerating tail base, and skeletal muscle [18].

## Availability of supporting data

Novel RNA-Seq data for *Sceloporus undulatus* samples are available under the NCBI accession identifiers listed in Table 1, and are associated with BioProject PRJNA371829. RNA-Seq data for the liver sample [23] were downloaded from NCBI from BioProject PRJNA183121, Run SRR629640. Datasets referring to the assembled and annotated transcriptome are available for download at Harvard Dataverse (doi:10.7910/DVN/EGRBCT).

### List of abbreviations

BLAST: Basic local alignment search tool
BUSCO: Benchmarking Universal Single-Copy Orthologs
GO: Gene Ontology
NCBI: National Center for Biotechnology Information
ORFs: open reading frames
RIN: RNA integrity number

## Authors’ contributions

MBG performed transcript assemblies and bioinformatics analyses; JJR performed transcript assemblies and bioinformatics analyses; MAT, OL, MJA and KK conceived the study; MAT and KK supervised bioinformatics analyses; MJA and OL provided samples. MBG drafted the manuscript, with edits from MT, KK, and MJA. All authors read and approved the final version.

## Acknowledgements

We thank Michael W. Sears and Colton D. Smith for collecting specimens and John Cornelius, Greer Dolby, Shawn Rupp, Timothy Webster and Cindy Xu for helpful discussions.

## Supporting information – Tables

**Table S1.** Contig length statistics for *Sceloporus undulatus de novo* assemblies.

|  | 1 tissue | 3 tissues | 4 tissues |
| --- | --- | --- | --- |
| Minimum length | 201.0 | 201.0 | 201.0 |
| 1<sup>st</sup> Quartile | 266.0 | 266.0 | 266.0 |
| Median | 382.0 | 377.0 | 375.0 |
| Mean | 829.9 | 822.4 | 781.0 |
| 3<sup>rd</sup> Quartile | 808.0 | 732.0 | 711.0 |
| Maximum length | 16,776.0 | 30,410.0 | 30,258.0 |

The Sund\_v1.0 assembly includes 4 tissues, specifically 3 tissues sequenced in this study (whole brain, skeletal muscle and embryos) and 1 previously reported tissue (liver [23]).

**Table S2.** Reads mapped to *Sceloporus undulatus de novo* Sund_v1.0 assembly.

| Read classification | Counts | Percentage of mapped reads |
| --- | --- | --- |
| Proper pairing | 170,981,981 | 97.10% |
| Left read only | 3,778,790 | 2.15% |
| Right read only | 1,015,874 | 0.58% |
| Improper pairing | 310,142 | 0.18% |

**Table S3.** Representation of full-length reconstructed protein-coding genes in *Sceloporus undulatus de novo* Sund_v1.0 transcriptome assembly, using the protein set of *Anolis carolinensis* (AnoCar2.0, Ensembl) as a reference.

| Alignment coverage | Counts | Cumulative counts |
| --- | --- | --- |
| 100% | 9,874 | 9,874 |
| 90% | 1,349 | 11,223 |
| 80% | 799 | 12,022 |
| 70% | 757 | 12,779 |
| 60% | 725 | 13,504 |
| 50% | 577 | 14,081 |
| 40% | 463 | 14,544 |
| 30% | 455 | 14,999 |
| 20% | 358 | 15,357 |
| 10% | 97 | 15,454 |

### Supporting information – Figure

**Figure S1.**
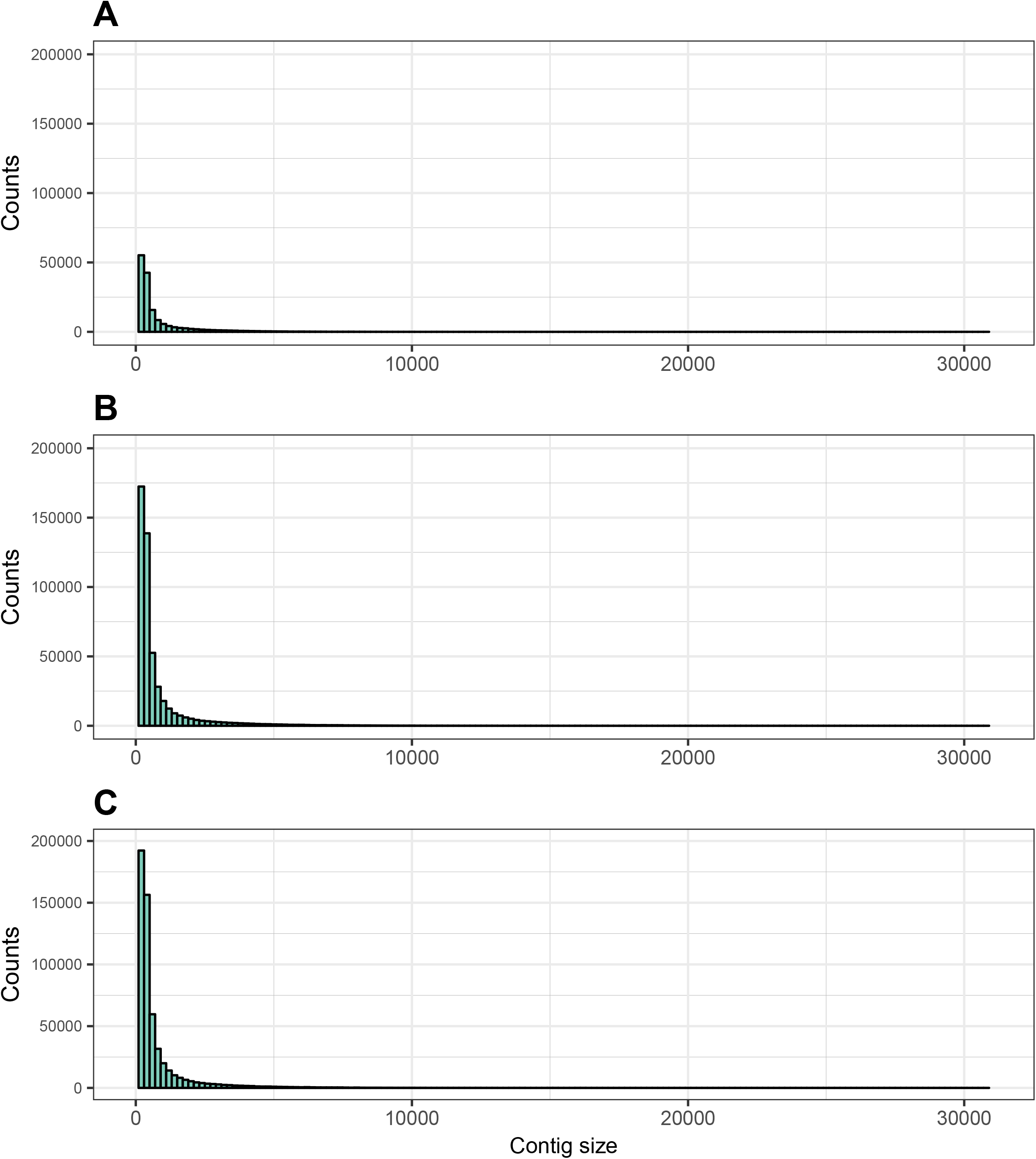
Contig sizes for different *Sceloporus undulatus* assemblies. Assemblies used **(A)** the previously published single tissue transcriptome (liver [23]), **(B)** transcriptomes from the 3 tissues sequenced in this study (brain, skeletal muscle and embryos), and **(C)** the combined data set of 4 tissues ([23] and this study).

